# Chromosome-scale *de novo* assembly and phasing of a Chinese indigenous pig genome

**DOI:** 10.1101/770958

**Authors:** Yalan Yang, Jinmin Lian, Bingkun Xie, Muya Chen, Yongchao Niu, Qiaowei Li, Yuwen Liu, Guoqiang Yi, Xinhao Fan, Yijie Tang, Jiang Li, Ivan Liachko, Shawn T. Sullivan, Bradley Nelson, Erwei Zuo, Zhonglin Tang

**Author notes:** These authors contributed equally to this work (Yalan Yang, Jinmin Lian, Bingkun Xie). To whom correspondence should be addressed. (Z.T.).

## Abstract

Chinese indigenous pigs differ significantly from Western commercial pig breeds in phenotypic and genomic characteristics. Thus, building a high-quality reference genome for Chinese indigenous pigs is pivotal to exploring gene function, genome evolution and improving genetic breeding in pigs. Here, we report an ultrahigh-quality phased chromosome-scale genome assembly for a male Luchuan pig, a representative Chinese domestic breed, by generating and combining data from PacBio Sequel reads, Illumina paired-end reads, high-throughput chromatin conformation capture and BioNano optical map. The primary assembly is ∼ 2.58 Gb in size with contig and scaffold N50s of 18.03 Mb and 140.09 Mb, respectively. Comparison between primary assembly and alternative haplotig reveals numerous haplotype-specific alleles, which provide a rich resource to study the allele-specific expression, epigenetic regulation, genome structure and evolution of pigs. Gene enrichment analysis indicates that the Luchuan-specific genes are predominantly enriched in Gene Ontology terms for phosphoprotein phosphatase activity, signaling receptor activity and phosphatidylinositol binding. We provide clear molecular evolutionary evidence that the divergence time between Luchuan and Duroc pigs is dated back to about 1.7 million years ago. Meanwhile, Luchuan exhibits fewer events of gene family expansion and stronger gene family contraction than Duroc. The positively selected genes (PSGs) in Luchuan pig significantly enrich for protein tyrosine kinase activity, microtubule motor activity, GTPase activator activity and ubiquitin-protein transferase activity, whereas the PSGs in Duroc pig enrich for G-protein coupled receptor activity. Overall, our findings not only provide key benchmark data for the pig genetics community, but also pave a new avenue for utilizing porcine biomedical models to study human health and diseases.

## Introduction

*Sus scrofa* (pig) is one of the most important domesticated animals for its enormous value in food supply and biomedical research. Plenty of archaeological and molecular evidence suggests that pigs were independently domesticated in the Near East and China about 9,000 years ago [1–3]. The effects of geographical divergence, local adaptation and artificial selection result in great phenotypic and genomic diversity among pigs from distinct locations and breeds [4, 5]. In China, there are ∼ 100 native breeds (China National Commission of Animal Genetic Resources 2011), accounting for about one-third of world breeds. To study pig genetics, the present pig reference genome (Sscrofa11.1) was derived from a Western pig (the Duroc breed) [6, 7]. However, Eastern and Western pigs have different genetic backgrounds. To better explore gene function, genome evolution and improve genetic breeding in pigs, it is of great value to build a reference genome for Chinese indigenous pigs.

Two main challenges for assembling a state-of-the-art high-quality reference genome are chromosome-scale contiguity and diploid phasing. Previous studies reported multiple *de novo* assemblies of Chinese native breeds using whole-genome shotgun-based strategies, and shed light on genomic and phenotype diversities of Chinese domestic pigs [4, 8–10]. Nonetheless, these shotgun-based approaches cannot yield large continuous genome scaffolds, significantly limiting the quality and contiguity of the current Chinese pig genome assemblies. Beyond genome assembly at the chromosome scale, accurate representation of haplotypes is crucial to identifying single-nucleotide polymorphisms (SNPs) and structural variants (SVs), haplotype structure and heterozygosities between two homologous chromosomes. Therefore, a phased genome assembly is essential for studies on intraspecific variation, allele-specific expression, epigenetic regulation, and chromosome evolution, as well as understanding how combinations of variants impact phenotypes [15–17]. Among the new technologies to tackle the two challenges in genome assembly, long-read sequencing, high-throughput chromatin conformation capture (Hi-C) and optical mapping technologies have been developed for ordering and orienting assembly contigs, and thus can create phased chromosome-scale genome assemblies [11]. These technologies have substantially improved genome assemblies for human, goat and gorilla [12–14]. However, a phased genome assembly with chromosome-scale contiguity for pigs is not yet to available, which results in the lack of resolution for pigs inter-haplotype variations, and impedes the dissection of the genetic basis of phenotypic differences in domestication between Eastern and Western pigs.

Here, we applied long-read sequencing (Pacbio), short paired-end reads (Illumina), Hi-C and optical map (BioNano) technologies to generate an assembly of the Luchuan pig, an indigenous breed from Guangxi province in South China. As a representative of the native breeds in China, Luchuan pig has many distinguishing phenotypic features comparing with Western domesticated pigs, including low growth rate, high fat content, excellent meat quality, early maturity, high fecundity, good maternal stability, wide adaptability to coarse feeding and strong disease resistance [4, 5]. To study the genetic basis underlying these phenotypic differences, Luchuan is an ideal material for building a high-quality reference genome representing Chinese indigenous pigs. In our study, a high-contiguous, chromosome-scale phased assembly of the Luchuan pig genome was *de novo* assembled. To our knowledge, this is the first published phased chromosome-scale assembly for mammals, providing important genetic resources and methodological references for future studies of animal genomic evolution, molecular breeding and biomedical research.

## Material and Methods

### Sample collection and sequencing

A Luchuan boar was obtained from the Institute of Animal Science of Guangxi province, China, for genome assembly. Genomic DNA was extracted from its blood sample. In order to generate a chromosome-scale assembly, four different genome libraries were constructed and sequenced according to the manufacturers’ instructions: (i) Whole genome sequencing (WGS) by PacBio Sequel platform (20-kb library); (ii) Hi-C chromosome conformation captured reads sequencing by Phase genomics; (iii) Short reads paired-end sequencing (150bp in length) by Illumina NovaSeq 6000 platform; (iv) BioNano optical map data (Nt.BspQI, Nb.BssSI and DLE-1 enzymes).

To fully assist genome annotation, thirty-seven RNAs from 14 tissues (heart, lung, adipose, kidney, liver, brain, spleen, stomach, leg muscle, dorsal muscles, testis, ovary, large intestine, small intestine) at four developmental stages (Days 0, 14, 50 and adult pigs) of Luchuan pigs (4 individuals) were equally pooled together. Two strand-specific RNA-seq libraries with an insert size of 350 bp using the NEBNext^®^ Ultra™ Directional RNA Library Prep Kit for Illumina^®^ (NEB, USA) were prepared and sequenced on an Illumina NovaSeq 6000 platform, to generate 150bp paired-end reads (Berry Genomics Co., Ltd., Tianjin, China). A PacBio full-length transcriptome library was constructed and sequenced on the Pacific Bioscience RS II sequencer (Berry Genomics, Co., Ltd., Beijing, China).

All animals and samples used in this study were collected according to the guidelines for the care and use of experimental animals established by the Ministry of Agriculture and Rural Affairs of China.

### *De novo* genome assembly and scaffolding

The primary contigs were assembled with the Falcon software packages (v2.0.5) [16] followed by the FALCON-Unzip and Arrow (v2.2.2) polishing, then a Hi-C-based contigs phasing was processed by FALCON-Phase to create phased, diploid contigs. Phase Genomics’ Proximo Hi-C genome scaffolding platform was used to establish chromosome-scale scaffolds from the draft assembly using a method similar to that described previously [14]. Following diploid chromosomal scaffolding, a round of polishing using Juicebox (v1.8.8) [18, 19] was performed to correct small errors in chromosome assignment, ordering and orientation. After a draft set of scaffolds was generated, FALCON-Phase was run again for Hi-C based scaffold phasing. The Illumina sequencing data were further used to improve the assembly by Pilon (v1.22) software. Given the availability of a relatively good quality of the Duroc pig (Sscrofa11.1) genome, a reference-assisted scaffolding strategy was conducted to get chromosome-level pseudomolecules with Chromosomer software (v0.1.4a) [20]. Quality control on the integrity of the assembly of genic regions was performed by using the independent BUSCO v3 benchmark (http://busco.ezlab.org/) [21].

### Assembly quality assessment

BioNano optical map data was used to assess the assembly quality, which produces physical maps with unique sequence motifs that can provide long-range structural information of the genome. Briefly, high-molecular weight DNA was extracted from the pig blood sample and digested with nickases Nt.BspQI, Nb.BssSI and Direct Labeling Enzyme 1 (DLE-1), respectively. After labeling and staining, DNA was loaded onto the Saphyr chip for sequencing. Raw data for each enzyme library were collected and converted into a BNX file by AutoDetect software, to obtain basic labeling and DNA length information. The filtered raw DNA molecules in BNX format were aligned, clustered and assembled into the BNG map by using the Bionano Solve pipeline. Two enzyme (Nt.BspQI, Nb.BssSI) hybrid scaffolding was firstly processed to produce a set of initial hybrid scaffold, a second round of hybrid scaffolding with genome map of DEL-1 enzyme was followed.

### Repeat annotation

There are two main types of repeats in the genome: tandem and interspersed. Tandem repetitive sequences were identified using Tandem Repeats Finder (TRF, version4.07). The interspersed repeat contents were identified using two methods: *de novo* repeat identification and known repeat searching against existing databases. RepeatModeler (version 1.0.8, http://www.repeatmasker.org/RepeatModeler/) was used to predict repeat sequences in the genome, and RepeatMasker (version 4.0.7) [22] was then used to search the Luchuan pig genome against the *de novo* transposable elements (TE) library. The homology-based approach involved applying commonly used databases of known repetitive sequences, RepeatMasker (version 4.0.7) and the Repbase database (version 21) [23] were used to identify TEsin the assembled genome. RepeatMasker and Repeat Protein Masker (http://repeatmasker.org) were applied for TEs identification at the DNA and protein levels, respectively.

### Gene prediction and annotation

Protein-coding region identification and gene prediction were conducted through a combination of three approaches as following:

i. Homology-based prediction. Protein sequences for human and five animal genomes (mouse, cattle, dog, goat and the Duroc pig) were downloaded from Ensembl release-95, and aligned to the Luchuan assembly using the TBLASTN program available in the BLAST v2.2.24 (E-value cutoff 1e-05). Then the SOLAR (version0.9.6), a dynamic program algorithm to link putative exons together, was employed to analyze the TBLASTN results. GeneWise (version 2.4.1) [24] was used to predict the exact gene structure of the corresponding genomic regions on each matched sequences;
ii. *De novo* prediction. Four *ab initio* gene prediction programs including Augustus (version 3.2.1) [25], GlimmerHMM (version 3.0.4) [26], Geneid (version 1.4.4) [27] and SNAP (version 2006-07-28) [24], were employed to predict coding regions in the repeat-masked genome;
iii. Transcriptome-based prediction methods. RNA-seq data (26.35 Gb) reads were mapped to the assembly using Hisat2 (version 2.1.0) [28]. Stringtie (version 1.2.2) and TransDecoder (version 3.0.1) were used to assemble the transcripts and identify candidate coding regions into gene models. For PacBio full-length transcriptome data (Iso-Seq), transcripts were identified by IsoSeq3 (version 3.1.0) with default parameters, then the Iso-Seq data were mapped to the reference genome with minimap2 (version 2.15-r905). Furthermore, Cupcake ToFU (v5.8) was used to get the final unique, full-length and high-quality isoforms of Pacbio data.

All gene models predicted based on the above three approaches were combined by EvidenceModeler (EVM) into a non-redundant set of gene structures, and the produced gene models were finally refined using the Program to Assemble Spliced Alignments (PASA v2.3.3) [29]. Functional annotation of protein-coding genes (PCGs) was achieved using BLASTP (E-value 1e-05) against two integrated protein sequence databases: SwissProt and TrEMBL. Protein domains were annotated by InterProScan (v5.30). The Gene Ontology (GO) terms for each gene were extracted with InterProScan [30]. The pathways in which the genes might be involved were assigned by BLAST against the KEGG databases (release 59.3) [31] with an E-value cutoff of 1e-05.

### Noncoding RNAs annotation

The transfer RNAs (tRNA) genes were predicted by tRNAscan-SE (version 1.3.1) [32] with eukaryote parameters. The ribosomal RNA (rRNA) fragments were predicted by aligning to human template rRNA sequences using BlastN (version 2.2.26) at an E-value of 1e-5. The microRNAs (miRNAs) and small nuclear RNAs (snRNAs) were detected by searching against the Rfam database (release 12.0) [33] with INFERNAL (version 1.1.1) [34]. Long non-coding RNAs (LncRNAs) and Circular RNAs (circRNAs) were predicted by methods described previously [4, 35, 36].

### Identification of orthologous gene sets across species

A gene family indicates a set of similar genes that descended from a single original gene in the last common ancestor of considered species. Orthologous gene sets of Luchuan pig, Duroc pig, cattle, goat, dog, mouse and human were used for genome comparisons. For a gene with multiple isoforms, we chose the longest transcript (≥ 50 amino acids) to represent the gene. The TreeFam methodology [37] was used to define a gene family and result in 3,733 single-copy orthologous genes for the six mammalian species. In addition, the one-to-one orthologous between these species were defined using BLASTP based on the Bidirectional Best Hit (BBH) method with a sequence coverage > 80% and identity > 80%, followed by selection of the best match.

### Variants calling

The primary assembly of Luchuan genome was aligned with the alternative haplotig assembly and the Duroc contigs by MUMmer (version 3.23)[38] with default parameters, and one-to-one genomic alignment results were extracted with the ‘delta-filter −1’ parameter. SNPs and indels were identified by show-snp from the one-to-one alignment blocks (parameter ‘-ClrT –x 1’). Structural variations were identified by Assemblytics (v1.0) software [39] base on the alignment blocks from MUMmer.

### Phylogenetic tree construction and evolution rate estimation

Single-copy gene families were used to construct a phylogenetic tree for Luchuan pig and the other mammalian genomes (Duroc pig, cattle, goat, dog, mouse and human). Four-fold degenerate sites were extracted from each family and concatenated into one supergene for each species. PhyML v3.0 was adopted to reconstruct the phylogenetic tree based on the GTR+gamma substitution model [40]. The divergence time among Luchuan pig, Duroc pig, cattle, goat, dog, mouse and human were estimated using the MCMCtree program (version 4.4) as implemented in the Phylogenetic Analysis of Maximum Likelihood (PAML) package [41], with an independent rates clock and HKY85 nucleotide substitution model. The calibration times (differentiation time between human and mouse, human and goat, cattle and goat, pig and goat) were derived from the TimeTree database [42].

## Results

### Assembly and phasing of the Luchuan pig genome

To construct a high-quality reference genome for Chinese indigenous pigs, a male Luchuan pig was used for WGS, which generated ∼140× Pacbio Sequel long reads (348.71 Gb), ∼41× Hi-C reads (102.42 Gb, Phase Genomics), ∼86× Illumina paired-end reads (214.48 Gb), and ∼351× BioNano optical map data (879.44 Gb, Bionano Genomics).

The Pacbio reads were first assembled *de novo*, producing an initial contig assembly with N50 of 18.68 Mb and a total length of 2.52 Gb. Then the assembly was integrated with Hi-C data to create phased diploid chromosome-scale scaffolds (Supplementary Figure 1), generating an alternative haplotype sequence with contig N50 of 18.79Mb, scaffold N50 of 141.24Mb and a total length of 2.55 Gb. After improving the assembly based on Illumina sequencing data, the optical map data were used to validate, correct and merge the scaffolds (Supplementary Table 1). Given the high quality of the present Duroc reference genome (Sscrofa11.1), a reference-assisted scaffolding strategy was used to get chromosome-level pseudomolecules (Figure 1A). Finally, we generated a high-contiguous, chromosome-scale and phased assembly of the Luchuan genome, yielding a 2.58 Gb primary assembly with a contig N50 of 18.03 Mb and a scaffold N50 of 140.09 Mb. This assembly is comparable in quality to the Duroc genome [7] and much better than other published pig genomes [4, 8–10] (Table 1). Remarkably, the alternative haplotig assembly size is very close to the primary assembly with a contig N50 of 17.77 Mb and a scaffold N50 of 140.08 Mb.

**Figure 1.**
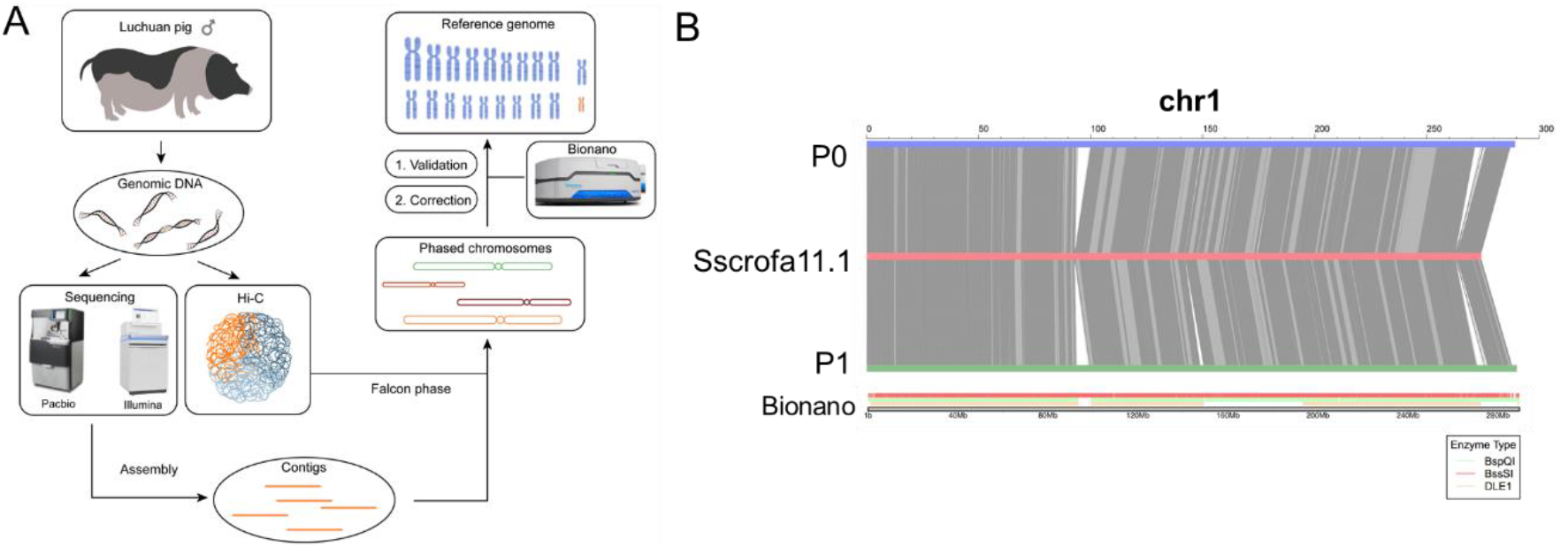
Genome Assembly. (A) The flowchart of contig, scaffold and chromosome assembly in this study. (B) Collinearity analysis for Chr1 between Sscrofa11.1 (*Middle*) and primary assembly (P0, *Upper*) and alternate haplotigs (P1, *Lower*) assemblies. Gray lines indicate collinearity between the genomes. Bionano optical map of Chr1 is shown in the bottom. Collinearity Analysis for other chromosomes were shown in Supplementary Figure 2.

**Table 1.** Comparison of features between the Luchuan pig and other assemblies.

|  | **Luchuan** | **Duroc<sup>[7]</sup>** | **Tibetan wild<sup>[8]</sup>** | **Wuzhishan<sup>[9]</sup>** | **Bama<sup>[10]</sup>** |
| --- | --- | --- | --- | --- | --- |
| **Sequenced genome size (Gb)** | 2.58 | 2.50 | 2.43 | 2.64 | 2.49 |
| **Contig N50 (Mb)** | 18.03 | 41.89 | 0.0207 | 0.0235 | 1.01 |
| **Scaffold N50 (Mb)** | 140.09 | 138.97 | 1.06 | 5.43 | 140.44 |
| **Percentage of anchoring and ordering** | 96.1% | 97.34% | - | - | 97.49% |
| **Predicted PCGs** | 22,710 | 22,452 | 21,806 | 20,326 | 21,334 |
| **Repeat proportion (%)** | 40.16 | 40.55 | 39.47 | 38.20 | 37.32 |
| **Complete BUSCOs (%)** | 95.1 | 96.0 | 93.1 | 95.2 | 93.9 |

\* Statistic of Duroc pig genome was based on Sscrofa11.1 (Ensembl release-95).

The reference assessment revealed that approximately 96.1% of the 2.58 Gb assembled final Luchuan assembly was assigned to 20 chromosomes (18 autosomes and X/Y chromosome) (Supplementary Table 2-3). The 20 chromosomes were made up of 466 contigs, reflecting the low fragmentation of these assemblies. We further evaluated the genome assembly quality, and found that 95.1% of the 4,104 core genes in the OrthoDB mammalian database were identified in the Luchuan primary assembly, of which 94.4% were single-copy, 0.7% duplicated, 2.9% fragmented and 2.0% missing (Supplementary Table 4).

### Validation of the phased diploid assemblies

The pseudo-chromosomes of Luchuan pig presented great colinearity with Sscrofa11.1, supporting a high-quality genome assembly (Figure 1B; Supplementary Figure 2). It is worth noting that the alternative haplotig also has highly collinear relationships with Duroc pig assembly (Supplementary Figure 2). Additionally, to assess the scaffolding accuracy of Luchuan assembly, we adopted the nickases Nt.BspQI, Nb.BssSI and DLE-1 for optical map library construction, and got 453 Gb, 345 Gb, and 618 Gb raw data using these three enzymes, respectively. After removing molecules in lengths less than 150 kb, we obtained 303 Gb, 268 Gb and 308 Gb high-quality optical molecules, accounting for > 100× coverage of genome size. The N50 of the molecules are 358 kb, 394 kb and 248 kb for Nt.BspQI, Nb.BssSI nickase and DLE-1 enzymes, respectively (Supplementary Table 5). The high concordance between the assembly and the optical map data provides strong support for the robustness of the assembly (Figure 2). By comparison between the contigs/scaffolds and optical maps, 74 and 73 conflicts were detected for the primary assembly and alternative haplotig, respectively. After conflict correction, we assembled 63 and 64 hybrid scaffolds based on genome map hybrid assembly for the primary assembly and alternative haplotig, respectively. These results demonstrated the high reliability of the alternate haplotype assembly.

**Figure 2.**
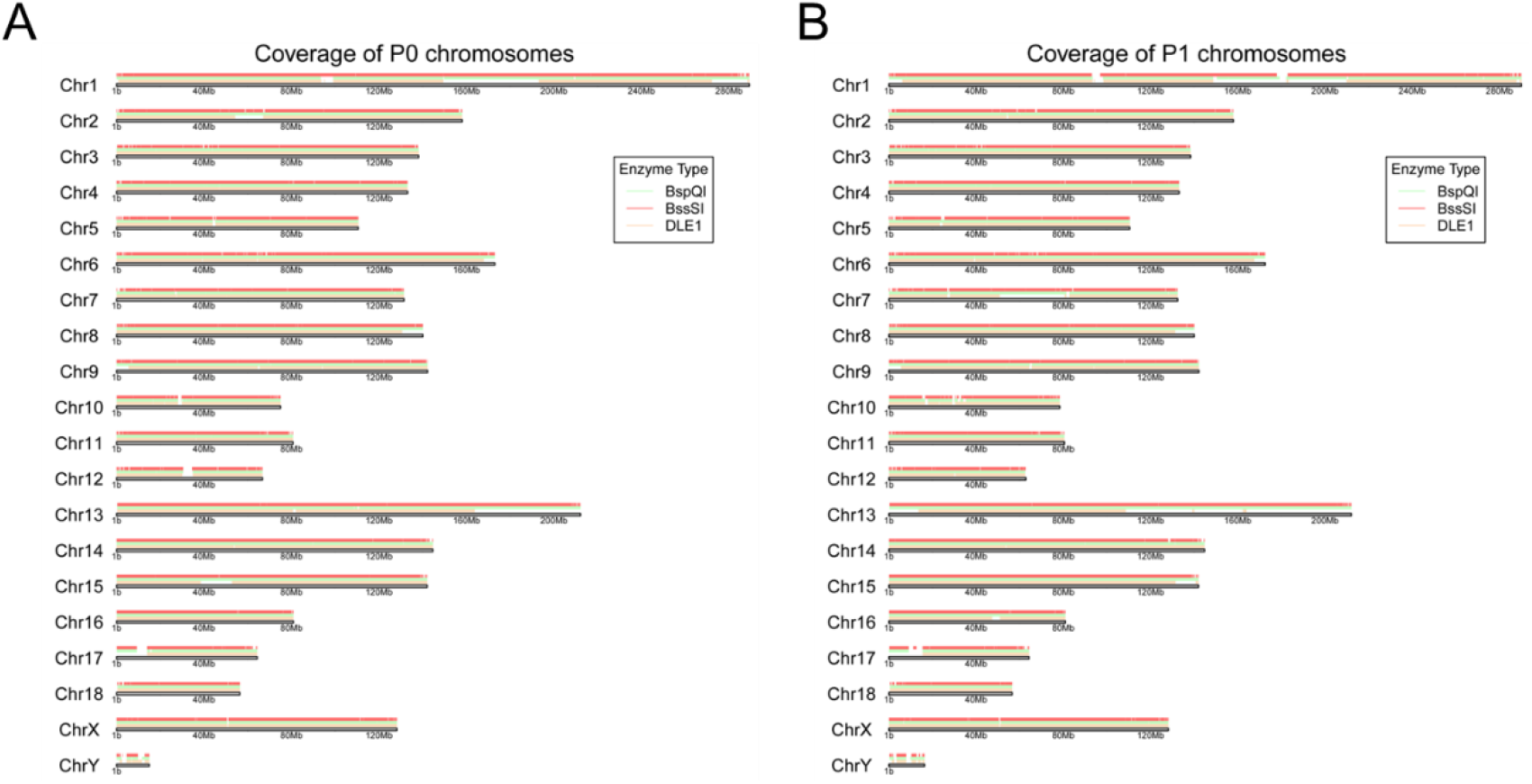
Assembly quality assessment by BioNano optical map data. (A) A comparison of Bionano optical maps and primary assembly of Luchuan pig. (B) A comparison of Bionano optical maps and alternate haplotigs of Luchuan pig. The optical genome maps are constructed by three enzymes (Nt.BspQI, Nb.BssSI and DLE-1) and shown in different colors. The black bar corresponds to the pseudo-chromosomes of Luchuan pig.

### Genetic variations between primary assembly and alternate haplotig

By comparing the primary assembly to the alternate haplotig, we identified numerous haplotype-specific alleles, including 6.83 million SNPs, 1.64 million short indels and 23,539 SVs (Figure 3). Among the SNPs, most (97.54%) were located in intergenic regions (63.01%) and intronic regions (34.51%), only 0.56% were located in coding sequences. Of the SNPs present in coding regions, 24,056 were synonymous and 13,959 were non-synonymous. In addition, 2,479 and 463 indels may result in frameshift and non-frameshift variations, respectively. These variations are valuable to further study the allele-specific expression, epigenetic regulation, genome structure and evolution in pigs.

**Figure 3.**
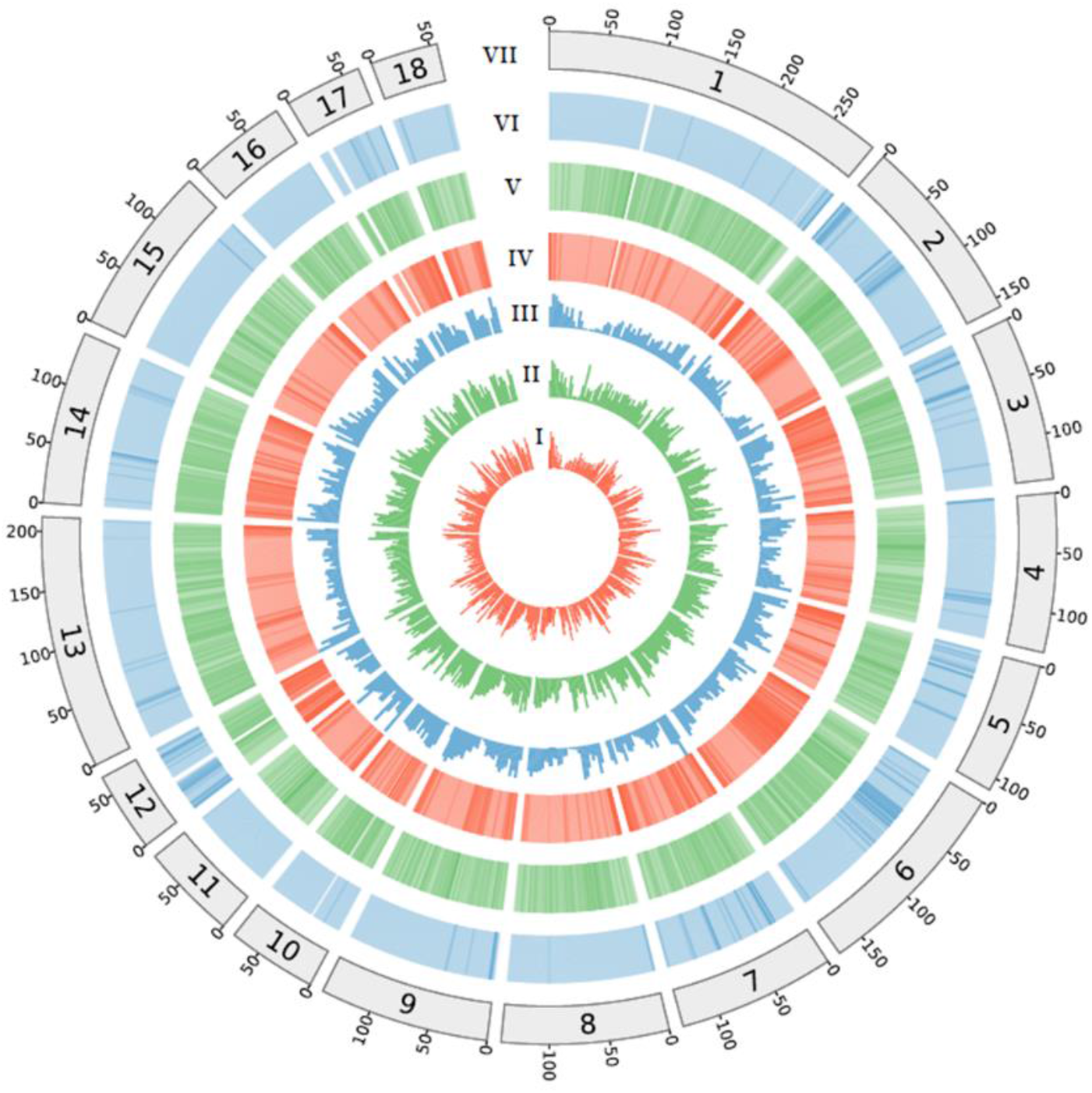
Circos plot showing the characterization of the Luchuan pig. I: Number of SNPs between primary assembly (P0) and alternate haplotigs (P1) in non-overlapping 5Mb windows; II: Number of indels between primary assembly (P0) and alternate haplotigs (P1) in non-overlapping 5Mb windows; III: Number of structural variants between primary assembly (P0) and alternate haplotigs (P1) in non-overlapping 5Mb windows; IV: GC content in non-overlapping 1Mb windows; V: Percent coverage of TEs in non-overlapping 1Mb windows; VI: Gene density calculated on the basis of the number of genes in non-overlapping 1Mb windows; VII: The length of pseudo-chromosome in the size of Mb.

### Genome annotation

We predicted a total of 22,710 PCGs with strong evidence in Luchuan by combining *ab initio* prediction, homologous protein prediction and transcriptome alignment. Of these PCGs, ∼90% gain clear supporting evidence based on transcriptome sequencing data and functional annotation information (Table 1; Supplementary Table 6). The average length of gene, exon and intron were 40,062bp, 177bp and 4,709bp, respectively. We also annotated 2,835 small ncRNAs including 388 miRNAs, 394 rRNAs, 1,076 tRNAs and 977 snRNAs. Additionally, 3,066 novel lncRNAs and 1,019 novel circRNAs were identified in the Luchuan genome (Supplementary Table 7).

Repeat elements accounted for ∼40.16% of the Luchuan genome (Supplementary Table 8-9). The two largest repeat classes were long-interspersed elements (LINEs) and short interspersed nuclear elements (SINEs), which comprised 27.83% and 10.86% of the genome, respectively. Tandem repeats constituted 3.88% of the genome. The number and length of genes and the proportion of repeat elements were similar to those present in the pig reference genome and other assemblies [4, 6–9].

### Comparative Genomic and Phylogenetic Analyses

We identified a total of 8,481 homologous gene families that are shared among Luchuan, Duroc, goat and human. Interestingly, there are 163 and 134 gene families specifically identified in Luchuan and Duroc, respectively (Figure 4A). Among those Luchan-specific gene families, 421 genes with supporting evidence of transcription or Interpro functional annotation were considered to be the high-quality Luchuan-specific genes. These genes are significantly (FDR < 0.05) enriched in GO terms for phosphoprotein phosphatase activity, signaling receptor activity and phosphatidylinositol binding. By comparison, the 207 Duroc-specific genes are functionally over-represented in biological processes related to actin filament binding, peptidase inhibitor activity, pheromone receptor activity, microtubule motor activity and epidermis development (Supplementary Table 10).

**Figure 4.**
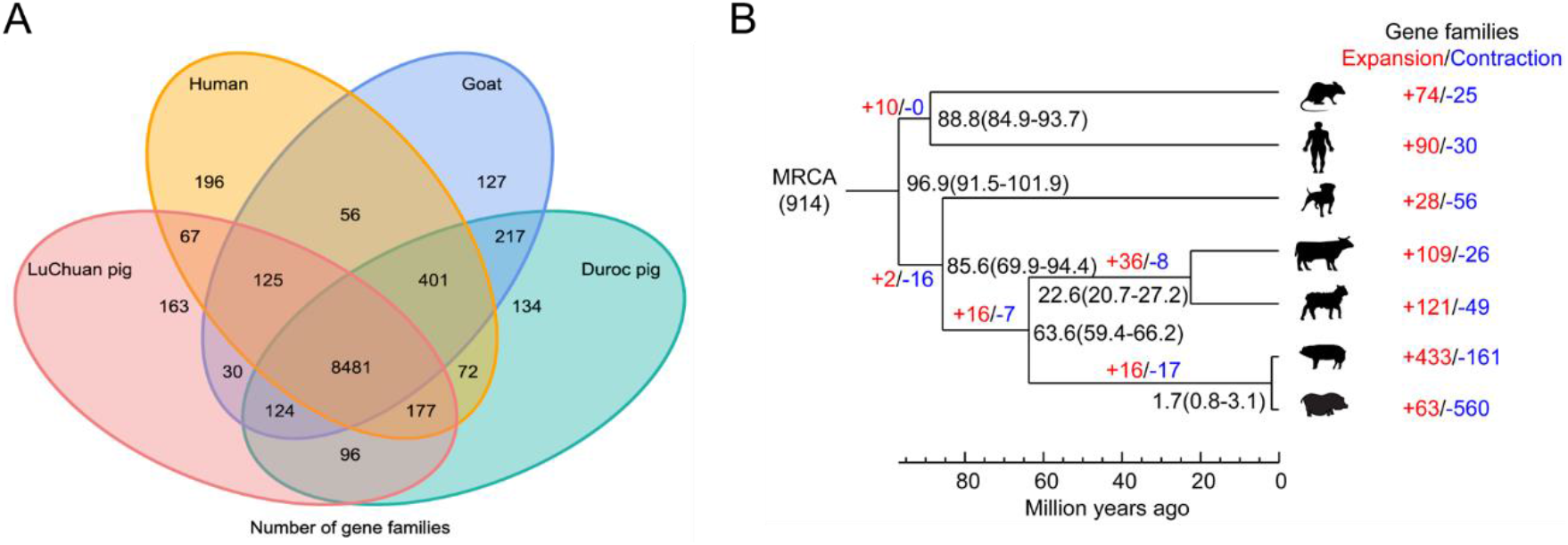
Comparative genomic and phylogenetic analyses. (A) Venn diagram showing shared orthologous gene families among genomes of Luchuan, Duroc, goat and human. (B) Phylogenetic tree with divergence times and history of orthologous gene families. Numbers on the nodes represent divergence times, with the error range shown in parentheses. The numbers of gene families that expanded (red) or contracted (blue) in each lineage after speciation are shown on the corresponding branch. MRCA, most recent common ancestor.

A phylogenetic tree was constructed using the pigs (Luchuan and Duroc) and five other mammals (cattle, goat, dog, human and mouse). As shown in Figure 4B, the divergence time between Luchuan and Duroc was estimated to be about 1.7 million years ago (MYA). Compared with Duroc, Luchuan showed fewer events of gene family expansion (63 vs. 433), and more events of gene family contraction (560 vs. 161) (Figure 4B). Notably, expanded genes in Luchuan were closely related to response to oxidative stress and biotic stimulus. In Duroc, the olfactory-related genes were significantly expanded, consistent with a previous study [8]. In addition, expanded genes in Duroc are significantly (P < 0.05) enriched in GO terms for galactosyltransferase activity, antioxidant activity and growth factor activity.

### Bidirectional selection between Luchuan and Duroc pigs

To study the bidirectional selection between Luchan and Duroc pigs, we further screened out 7,222 one-to-one orthologous gene sets from the seven mammals. We found 272 and 768 positively selected genes (PSGs) in the Luchuan and Duroc pigs (P < 0.05, likelihood ratio test), respectively. It is worth noting that 25 PSGs were shared in both breeds, such as *CACNA1F*, a calcium channel subunit gene, and *RBM46*, an RNA binding motif protein. Enrichment analysis revealed the PSGs detected in Luchuan were especially enriched in GO terms related to protein tyrosine kinase activity (8 PSGs), microtubule motor activity (6 PSGs), GTPase activator activity (6 PSGs) and ubiquitin-protein transferase activity (6 PSGs), whereas PSGs in Duroc pigs were significantly enriched in GO terms for G-protein coupled receptor activity, which is closely related with the olfactory receptors (Supplementary Table 11).

## Discussion

Chinese indigenous and Western pigs are independently domesticated and exhibit a great spectrum of phenotypic and genomic differences [4, 6–9]. A comprehensive exploration of the genetic diversity within and between pig breeds is important for animal breeding and biomedical research. The present pig reference genome (Sscrofa11.1) was derived from a Western breed (Duroc pig) [7] with high continuity and quality. Increased accessibility to short-read sequencing has resulted in a deluge of genome assemblies for Chinese indigenous pigs, although incomplete and fragmented compared with Duroc [4, 8–10]. Until now, no chromosome-level phased assemblies of Chinese indigenous pigs have been built, so accurately investigating the full range of genetic variations and phased diploid architecture is extremely difficult. Recently, rapid progress in high-throughput DNA sequencing and library preparation methods have enabled the generation of phased genome assemblies with chromosome-level quality [15–17]. Built upon these most recent technology breakthroughs, here we present, to our knowledge, the first phased chromosome-scale genome assembly of pigs, which is also the first such type of published assembly for mammals. Our genome assembly yields a 2.58 Gb primary assembly with a contig N50 of 18.03 Mb, with comparable quality to the current reference genome [7].

Synteny analysis revealed strong collinearity between the genomes of Luchuan and Duroc pigs, supporting great overall quality of our assembly. Notably, our assembly approach also makes it possible to construct a high-quality alternate haplotig assembly, which is comparable to the primary assembly with a scaffold N50 size of 17.77 Mb. Using the phased diploid assembly, we are able to identify structural variations between two homologous chromosomes [15, 16], which are important for understanding how combinations of variants impact phenotypes. Millions of genetic variations between primary assembly and alternate haplotig of Luchuan genome were identified in our study, which provided an unprecedentedly detailed resource to further study the allele-specific expression, epigenetic regulation, genome structure and evolution of Eastern pigs [15–17]. Moreover, combining our Luchuan genome and the classic Duroc assembly would provide foundational resources to study the genetic basis underlying the phenotypic differences between Eastern and Western pigs.

To study the evolution and domestication of Luchuan pigs, we reconstructed the phylogenetic tree among Luchuan, Duroc, cattle, goat, dog, human and mouse. Our analysis revealed that the divergence time between Luchuan and Duroc was about 1.7 MYA, which is in close proximity to the split time between Asian and European wild boars (0.8-2 MYA) [43–45]. Gene replication is one of the basic mechanisms for acquiring new functions and physiological features, and accordingly studying gene family expansion and contraction provides unique perspectives on the genetic basis of local domestication and adaptation [46, 47]. The Luchuan genome exhibited fewer events of gene family expansion and stronger gene family contraction compared with Duroc pig, which is in accordance with the comparative analysis between Duroc pig and Tibetan wild boar [8]. Duroc pig was reported to have markedly more olfactory-related genes than Tibetan wild boar [8]. Our results also confirmed that these genes were significantly expanded and positively selected in Duroc pig. The oxidative stress and response to biotic stimulus-related genes were expanded and GTPase activator activity-related genes were positively selected in Luchuan pig, which might confer the remarkable capabilities of Luchuan to adapt to coarse feeding and strong resistance to diseases, which are important features shared by many Chinese indigenous breeds. The PSGs analysis suggested that the Duroc pig had experienced stronger selection pressures during breeding than Luchuan pig. These results provided novel insights into the distinct evolutionary scenarios occurring under different local adaptation and artificial selection between Chinese indigenous and Western pig breeds.

Overall, we presented the first phased chromosome-scale genome assembly of a Chinese indigenous breed, which provides great resources for understanding pig evolution and domestication. This Luchuan pig genome assembly would benefit the dissection of the genetic basis and molecular mechanisms underlying phenotypic differences between and within pig breeds, facilitate molecular breeding to improve economical traits, and shed light on the etiology of human traits and diseases.

## Acknowledgements

This work was supported by National Natural Science Foundation of China (31830090), the National Key Project (2016ZX08009-003-006), the Shenzhen Dapeng New District Special Fund for Industry Development (KY20180114) and the Agricultural Science and Technology Innovation Program (ASTIP-AGIS5).

## Author contributions

Z.L.T conceived, coordinated and managed the project; Y.L.Y, Y.W.L, G.Q.Y, M.Y.C, Y.C.N, J.M.L, J.L, I.L, S.T.S and B.N assembled and annotated the genome sequences, and carried out other computational and bioinformatics analysis; B.K.X provided the Luchuan pigs and helped in samples collection. Z.L.T, X.H.F, Y.L.Y and Y.J.T performed animal experiment and collected biological samples; Y.L.Y, Y.C.N and J.M.L wrote the manuscript; Z.L.T, Y.L.Y, G.Q.Y, and E.W.Z revised the paper. All authors read and approved the final manuscript.

---

**Supplementary Figure 1.**
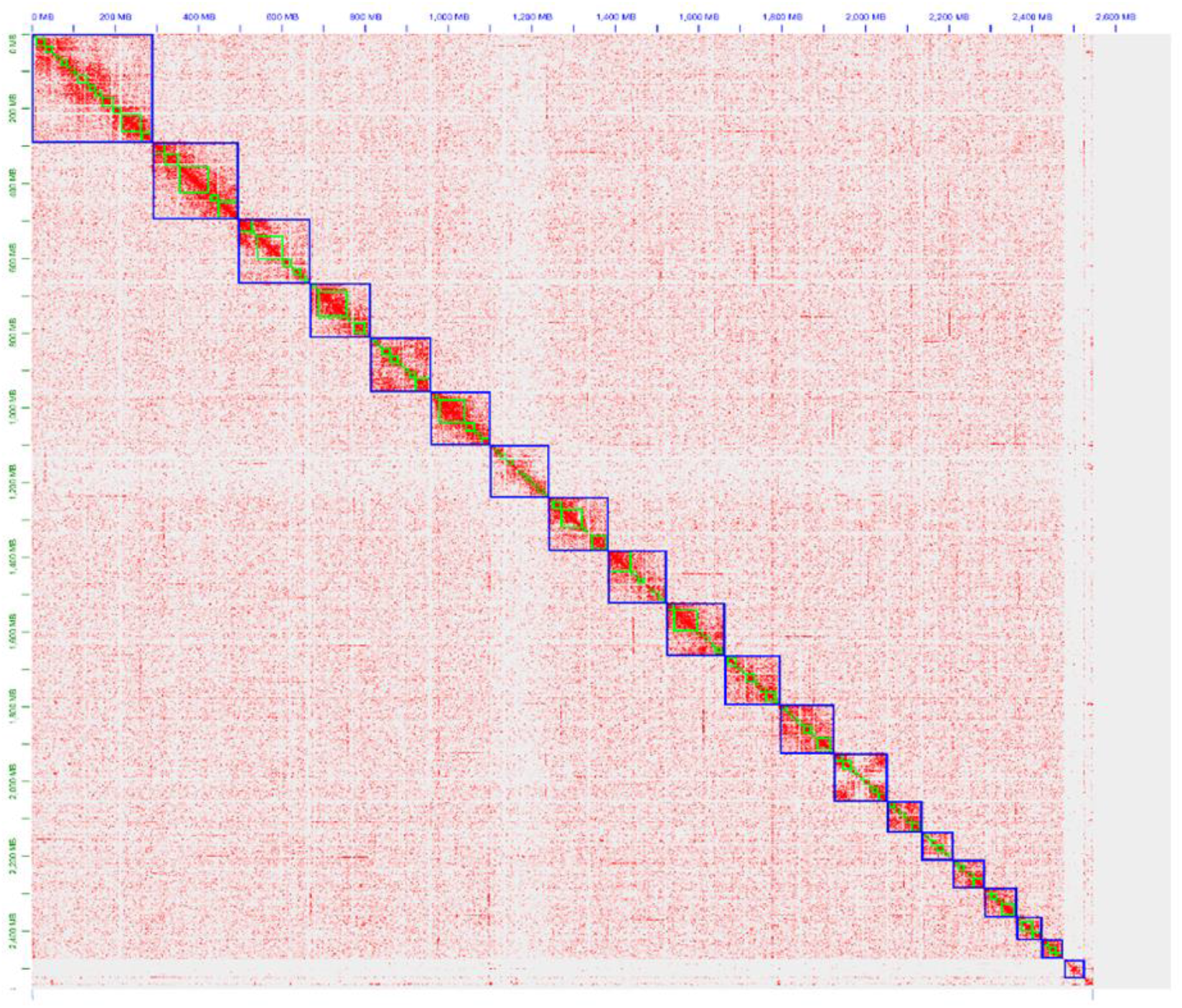
HiC contact heatmap. Genome-wide analysis of chromatin interactions in Luchuan genome.

**Supplementary Figure 2.**
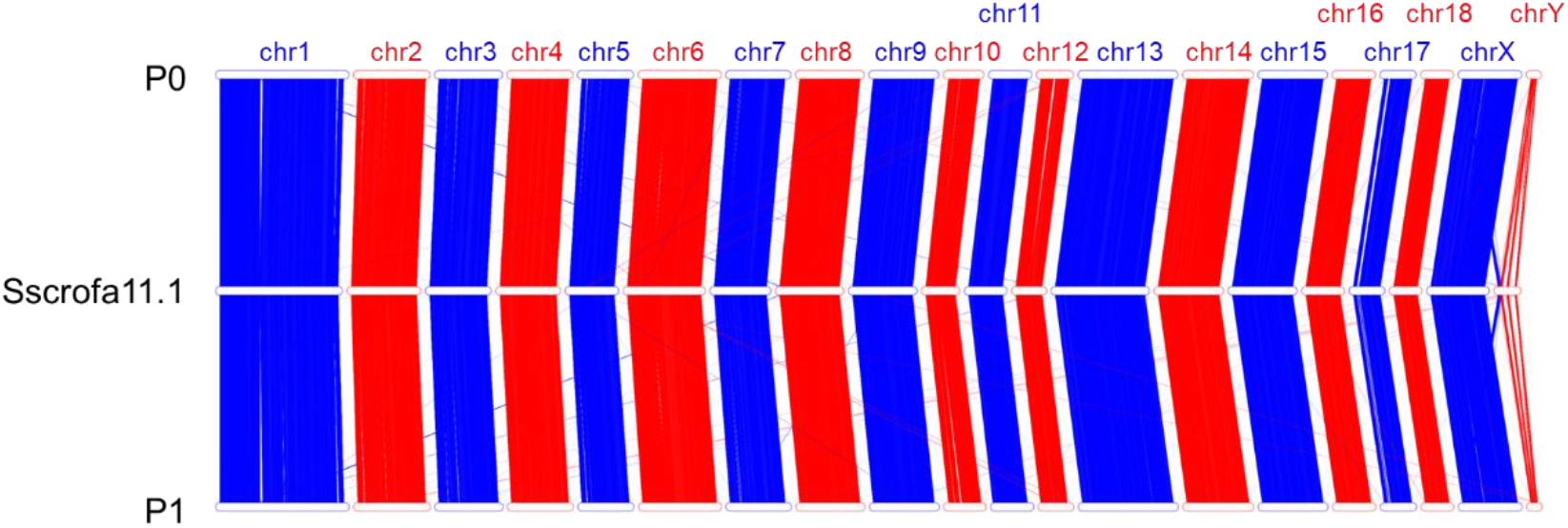
Collinearity analysis between Sscrofa11.1 (*Middle*) and primary assembly (P0, *Upper*) and alternate haplotigs (P1, *Lower*) assemblies. Red and blue lines indicate collinearity between the genomes.

