## Supplemental Tables for "Chromosome-scale *de novo* assembly and phasing of a Chinese indigenous pig genome"

**Supplementary Table 1. Statistics of the two haplotigs in each step of assembly processes.**

| | Primary assembly (P0) | | | Alternate haplotigs (P1) | | |
|---|---|---|---|---|---|---|
| | Total size (Gb) | Contig N50 (Mb) | Scaffold N50 (Mb) | Total size (Gb) | Contig N50 (Mb) | Scaffold N50 (Mb) |
| Falcon-unzip (Contig Phasing) | 2.519 | 18.667 | - | 1.902 | 0.426 | - |
| Falcon-phase (Hi-C based contig phasing) | 2.548 | 18.663 | - | 2.550 | 18.794 | - |
| Proximo + Falcon-phase (Hi-C based scaffold phasing) | 2.545 | 18.663 | 141.275 | 2.545 | 18.794 | 141.243 |
| Pilon (Indel Correction) | 2.544 | 18.559 | 141.216 | 2.544 | 18.554 | 141.173 |
| Bionano (Optical mapping based correction and scaffolding) | 2.544 | 18.025 | 77.560 | 2.543 | 17.774 | 77.553 |
| Chromosomer (reference assisted scaffolding) | 2.583 | 18.025 | 140.090 | 2.578 | 17.774 | 140.078 |

**Supplementary Table2. The final assembly of Luchuan pig.**

| | Primary assembly (P0) | | | | Alternate haplotigs (P1) | | | |
|---|---|---|---|---|---|---|---|---|
| | Contigs | | Super-Scaffolds (chromosome level) | | Contigs | | Super-Scaffolds (chromosome level) | |
| | Size (bp) | Number | Size (bp) | Number | Size (bp) | Number | Size (bp) | Number |
| **N90** | 3,137,147 | 159 | 66,690,848 | 17 | 3,138,126 | 163 | 64,260,881 | 17 |
| **N80** | 7,047,427 | 110 | 80,868,033 | 14 | 6,969,292 | 113 | 80,878,586 | 14 |
| **N70** | 10,379,127 | 80 | 128,274,157 | 12 | 10,478,028 | 82 | 132,461,062 | 11 |
| **N60** | 13,670,071 | 58 | 133,063,186 | 10 | 13,564,678 | 61 | 133,137,529 | 10 |
| **N50** | 18,025,403 | 42 | 140,090,008 | 8 | 17,774,162 | 44 | 140,077,844 | 8 |
| **Longest** | 71,316,358 | - | 289,388,999 | - | 64,286,259 | - | 290,116,104 | - |
| **Total Size** | 2,543,540,403 | - | 2,582,939,293 | - | 2,543,466,903 | - | 2,578,303,356 | - |
| **Number (>=100kb)** | - | 599 | - | 246 | - | 600 | - | 242 |
| **Number (>=1Mb)** | - | 217 | - | 28 | - | 221 | - | 27 |
| **Number (>=10Mb)** | - | 82 | - | 20 | - | 85 | - | 20 |
| **Number (>=50Mb)** | - | 5 | - | 19 | - | 4 | - | 19 |

**Supplementary Table 3. Statistic of pseudo-chromosomes length of primary assembly**

| pseudo-chromosome | Luchuan | Sscrofa11.1 (Duroc) |
|---|---|---|
| **1** | 289,388,999 | 274,330,532 |
| **2** | 157,994,446 | 151,935,994 |
| **3** | 138,093,187 | 132,848,913 |
| **4** | 133,063,186 | 130,910,915 |
| **5** | 110,581,696 | 104,526,007 |
| **6** | 172,988,264 | 170,843,587 |
| **7** | 131,499,371 | 121,844,099 |
| **8** | 140,090,008 | 138,966,237 |
| **9** | 142,256,968 | 139,512,083 |
| **10** | 74,986,007 | 69,359,453 |
| **11** | 80,691,476 | 79,169,978 |
| **12** | 66,690,848 | 61,602,749 |
| **13** | 212,125,878 | 208,334,590 |
| **14** | 144,624,675 | 141,755,446 |
| **15** | 142,062,807 | 140,412,725 |
| **16** | 80,868,033 | 79,944,280 |
| **17** | 64,381,442 | 63,494,081 |
| **18** | 56,486,588 | 55,982,971 |
| **X** | 128,274,157 | 125,939,595 |
| **Y** | 14,970,572 | 43,547,828 |
| **Un-anchored** | 100,820,685 | 66,650,325 |
| **Gap size** | 39,398,890 (1.53%) | 29,864,684 (1.19%) |
| **Total size** | 2,582,939,293 | 2,501,912,388 |

**Supplementary Table4. Comparison of the pig assemblies gene-space with the 4,104 BUSCO mammalian gene set.**

| | **Luchuan** | **Duroc** | **Tibetan wild** | **Wuzhishan** | **Bama** |
|---|---|---|---|---|---|
| Complete BUSCOs*(%) | 95.1 | 96.0 | 93.1 | 95.2 | 93.9 |
| Single copy (%) | 94.4 | 95.4 | 92.6 | 94.7 | 93.3 |
| Duplicated copy(%) | 0.7 | 0.6 | 0.5 | 0.5 | 0.6 |
| Fragmented (%) | 2.9 | 2.4 | 3.8 | 3.0 | 3.2 |
| Missing (%) | 2.0 | 1.6 | 3.1 | 1.8 | 2.9 |

BUSCOs analysis included 4,104 BUSCO mammalia gene set, BUSCO were run with the "--mode genome--limit 20 --long"parameters.

**Supplementary Table 5. BioNano optical maps of Luchuan pig**

<table>
<tr><th colspan="2">Enzyme</th><th>Molecules Number</th><th>TotalLength (Mb)</th><th>Molecule N50 (kb)</th><th>Average length (kb)</th><th>Label Density (/100kb)</th></tr>
<tr><td colspan="2">BspQI</td><td>931,439</td><td>303,138</td><td>358</td><td>325</td><td>9.600</td></tr>
<tr><td colspan="2">BssSI</td><td>756,382</td><td>268,280</td><td>394</td><td>354</td><td>10.488</td></tr>
<tr><td colspan="2">DLE-1</td><td>1,231,833</td><td>308,029</td><td>248</td><td>250</td><td>14.427</td></tr>
<tr><th colspan="7">Optical maps information</th></tr>
<tr><th></th><th>Enzyme</th><th>Total Genome Map Length (Mb)</th><th colspan="2">Avg. Genome Map Length (Mb)</th><th>Median Genome Map Length (Mb)</th><th>Genome Map N50(Mb)</th></tr>
<tr><td rowspan="3">Primary assembly (P0)</td><td>BspQI</td><td>2568.374</td><td colspan="2">3.167</td><td>2.14</td><td>5.037</td></tr>
<tr><td>BssSI</td><td>2548.967</td><td colspan="2">1.722</td><td>0.897</td><td>3.331</td></tr>
<tr><td>DLE-1</td><td>2523.792</td><td colspan="2">19.564</td><td>1.646</td><td>65.083</td></tr>
<tr><td rowspan="3">Alternate haplotigs (P1)</td><td>BspQI</td><td>2564.096</td><td colspan="2">3.205</td><td>2.079</td><td>5.112</td></tr>
<tr><td>BssSI</td><td>2544.498</td><td colspan="2">1.729</td><td>0.902</td><td>3.339</td></tr>
<tr><td>DLE-1</td><td>2526.098</td><td colspan="2">19.735</td><td>1.663</td><td>65.15</td></tr>
<tr><th colspan="7">Hybrid genome assembly</th></tr>
<tr><th></th><th>Hybrid strategy</th><th>Scaffold Length(bp)</th><th colspan="2">Scaffold Number</th><th>Max Scaffold Length(bp)</th><th>Scaffold N50 (bp)</th></tr>
<tr><td rowspan="2">Primary assembly (P0)</td><td>BssSI+ BspQI</td><td>2,575,271,988</td><td colspan="2">1,355</td><td>142,870,408</td><td>69,255,431</td></tr>
<tr><td>BssSI+ BspQI+ DLE-1</td><td>2,583,135,539</td><td colspan="2">1,337</td><td>212,125,878</td><td>77,560,082</td></tr>
<tr><td rowspan="2">Alternate haplotigs (P1)</td><td>BssSI+ BspQI</td><td>2,569,480,440</td><td colspan="2">1,361</td><td>212,144,969</td><td>77,553,416</td></tr>
<tr><td>BssSI+ BspQI+ DLE-1</td><td>2,578,218,356</td><td colspan="2">1,345</td><td>212,144,995</td><td>77,553,416</td></tr>
</table>

**Supplementary Table6. General statistics of predicted protein-coding genes in the Luchuan pig genome compared with other representative mammalian genomes.**

| Gene set | Num ber | Average gene length (bp) | Average mRNA length (bp) | Average exons per gene | Average exon length (bp) | Average intron length (bp) |
|---|---|---|---|---|---|---|
| **Luchuan** | 22,710 | 40,062 | 1,472 | 8.34 | 177 | 4,709 |
| **Duroc** | 22,452 | 42,152 | 1,515 | 8.80 | 172 | 4,543 |
| **Cattle** | 21,867 | 41,372 | 1,564 | 8.71 | 180 | 4,733 |
| **Goat** | 21,361 | 37,769 | 1,560 | 8.98 | 174 | 4,250 |
| **Dog** | 19,856 | 41,466 | 1,644 | 9.53 | 172 | 4,307 |
| **Human** | 23,358 | 63,260 | 1,699 | 9.75 | 174 | 5,505 |
| **Mouse** | 22,600 | 42,841 | 1,593 | 8.89 | 179 | 4,515 |

**Supplementary Table 7. Non-coding RNAs in the Luchuan pig assembly.**

| Type | | Number | Average length(bp) | Total length(bp) | % of genome |
|---|---|---|---|---|---|
| miRNA | | 388 | 84 | 32,520 | 0.001279 |
| tRNA | | 1,076 | 74 | 79,626 | 0.003131 |
| rRNA | rRNA | 394 | 93 | 36,596 | 0.001439 |
| | 18S | 27 | 298 | 8,042 | 0.000316 |
| | 28S | 115 | 105 | 12,052 | 0.000474 |
| | 5.8S | 3 | 76 | 227 | 0.000009 |
| | 5S | 249 | 65 | 16,275 | 0.00064 |
| snRNA | snRNA | 977 | 113 | 110,068 | 0.004327 |
| | CD-box | 215 | 88 | 18,860 | 0.000741 |
| | HACA-box | 210 | 137 | 28,696 | 0.001128 |
| | splicing | 530 | 112 | 59,123 | 0.002324 |
| LncRNA | | 3,066 | 921 | 2,823,510 | 0.111007 |
| circRNA | | 1,019 | 44,522 | 45,367,891 | 1.783651 |

**Note:** '% of genome' was calculated by the non-gap genome size 2,543,540,403 bp. microRNA (miRNA), small nuclear RNA (snRNA) and tRNA located in repeat or gap regions were filtered. rRNA with identity less than 85% were also filtered.

**Supplementary Table8. General statistics of repeats in the Luchuan pig genome.**

| Type | Repeat Size | % of genome |
|---|---|---|
| **Tandem Repeats** | 100,157,191 | 3.88 |
| **Interspersed repeat** | 961,475,196 | 37.22 |
| **Total** | 1,037,274,380 | 40.16 |

Note: Some elements may partly include another element domain.

**Supplementary Table9. TEs content in the assembled Luchuan pig genome.**

| Type | Repbase TEs | | TE protiens | | De novo | | Combined TEs* | |
|---|---|---|---|---|---|---|---|---|
| | Length (bp) | % in genome | Length (bp) | % in genome | Length (bp) | % in genome | Length (bp) | % in genome |
| **DNA** | 20,642,545 | 0.80 | 4,619,459 | 0.18 | 32,853,798 | 1.27 | 41,788,657 | 1.62 |
| **LINE** | 422,313,488 | 16.35 | 231,065,996 | 8.95 | 663,619,398 | 25.69 | 718,797,861 | 27.83 |
| **SINE** | 262,535,079 | 10.16 | - | 0.00 | 22,215,646 | 0.86 | 280,473,765 | 10.86 |
| **LTR** | 64,503,517 | 2.50 | 8,580,071 | 0.33 | 81,770,447 | 3.17 | 99,520,517 | 3.85 |
| **†Unknown** | - | 0.00 | - | 0.00 | 9,420,682 | 0.36 | 9,420,682 | 0.36 |
| **Total** | 807,514,294 | 31.26 | 244,224,272 | 9.46 | 805,510,909 | 31.19 | 961,475,196 | 37.22 |

Note: This statistical table does not contain Tandem Repeats, some elements may partly include another element domain.

*Combined: the non-redundant consensus of all repeat prediction/classification methods employed.

†Unknown: the predicted repeats that cannot be classified by RepeatMasker;

LINE, long interspersed nuclear elements; SINE, short interspersed nuclear elements; LTR, long terminal repeat.

**Supplementary Table 10. Functional gene categories enriched for the Luchuan pig and Duroc pig specific families.**

| GO_domain | GO_ID | Number | qvalue | GO_description |
|---|---|---|---|---|
| | | **Luchuan pig** | | |
| **component** | GO:0005634 | 10 | 0.012836 | nucleus |
| **function** | GO:0004721 | 8 | 6.64E-12 | phosphoprotein phosphatase activity |
| **function** | GO:0038023 | 4 | 0.002448 | signaling receptor activity |
| **function** | GO:0035091 | 3 | 0.014523 | phosphatidylinositol binding |
| **process** | GO:0006397 | 8 | 2.47E-09 | mRNA processing |
| **process** | GO:0015074 | 4 | 5.96E-06 | DNA integration |
| | | **Duroc pig** | | |
| **component** | GO:0005635 | 4 | 1.93E-06 | nuclear envelope |
| **component** | GO:0034993 | 4 | 2.22E-06 | meiotic nuclear membrane microtubule tethering complex |
| **component** | GO:0005576 | 6 | 0.038255 | extracellular region |
| **function** | GO:0051015 | 6 | 1.72E-06 | actin filament binding |
| **function** | GO:0030414 | 4 | 3.72E-05 | peptidase inhibitor activity |
| **function** | GO:0016503 | 3 | 0.000168 | pheromone receptor activity |
| **function** | GO:0003777 | 3 | 0.023113 | microtubule motor activity |
| **process** | GO:0008544 | 5 | 2.67E-08 | epidermis development |
| **process** | GO:0090286 | 4 | 1.72E-06 | cytoskeletal anchoring at nuclear membrane |
| **process** | GO:0007018 | 3 | 0.023113 | microtubule-based movement |

**Supplementary Table 11. Functional gene categories enriched for the PSGs of Luchuan pig and Duroc pig.**

| GO domain | GO ID | Number of genes | qvalue | GO_description |
|---|---|---|---|---|
| | | **PSGs of Luchuan pig** | | |
| **component** | GO:0030286 | 4 | 1.98E-02 | dynein complex |
| **function** | GO:0004713 | 8 | 7.26E-03 | protein tyrosine kinase activity |
| **function** | GO:0004714 | 5 | 7.26E-03 | transmembrane receptor protein tyrosine kinase activity |
| **function** | GO:0005515 | 65 | 7.26E-03 | protein binding |
| **function** | GO:0005524 | 32 | 1.10E-02 | ATP binding |
| **function** | GO:0003777 | 6 | 4.29E-02 | microtubule motor activity |
| **function** | GO:0005096 | 6 | 4.29E-02 | GTPase activator activity |
| **function** | GO:0004842 | 6 | 4.44E-02 | ubiquitin-protein transferase activity |
| **process** | GO:0007169 | 7 | 1.94E-03 | transmembrane receptor protein tyrosine kinase signaling pathway |
| | | **PSGs of Duroc pig** | | |
| **component** | GO:0016021 | 56 | 7.98E-07 | integral component of membrane |
| **function** | GO:0004930 | 8 | 9.32E-13 | G-protein coupled receptor activity |
| **function** | GO:0005524 | 79 | 2.97E-07 | ATP binding |
| **function** | GO:0005515 | 147 | 1.30E-04 | protein binding |
| **process** | GO:0007186 | 8 | 4.62E-14 | G-protein coupled receptor signaling pathway |
